# The complete mitochondrial genome of *Calyptogena marissinica* (Heterodonta: Veneroida: Vesicomyidae): insight into the deep-sea adaptive evolution of vesicomyids

**DOI:** 10.1101/648121

**Authors:** Mei Yang, Lin Gong, Jixing Sui, Xinzheng Li

## Abstract

The deep sea is one of the most extreme environments on earth, with low oxygen, high hydrostatic pressure and high levels of toxins. Species of the family Vesicomyidae are among the dominant chemosymbiotic bivalves found in this harsh habitat. Mitochondria play a vital role in oxygen usage and energy metabolism; thus, they may be under selection during the adaptive evolution of deep-sea vesicomyids. In this study, the mitochondrial genome (mitogenome) of the vesicomyid bivalve *Calyptogena marissinica* was sequenced with Illumina sequencing. The mitogenome of *C. marissinica* is 17,374 bp in length and contains 13 protein-coding genes, 2 ribosomal RNA genes (*rrnS* and *rrnL*) and 22 transfer RNA genes. All of these genes are encoded on the heavy strand. Some special elements, such as tandem repeat sequences, “G(A)_n_T” motifs and AT-rich sequences, were observed in the control region of the *C. marissinica* mitogenome, which is involved in the regulation of replication and transcription of the mitogenome and may be helpful in adjusting the mitochondrial energy metabolism of organisms to adapt to the deep-sea environment. The gene arrangement of protein-coding genes was identical to that of other sequenced vesicomyids. Phylogenetic analyses clustered *C. marissinica* with previously reported vesicomyid bivalves with high support values. Positive selection analysis revealed evidence of adaptive change in the mitogenome of Vesicomyidae. Ten potentially important adaptive residues were identified, which were located in *cox1, cox3, cob, nad2, nad4* and *nad5*. Overall, this study sheds light on the mitogenomic adaptation of vesicomyid bivalves that inhabit the deep-sea environment.

## Introduction

Mitochondria, which descended from proteobacteria via endosymbiosis, are important organelles in eukaryotic cells and are involved in various processes, such as ATP generation, signaling, cell differentiation, growth and apoptosis [1]. Moreover, mitochondria have their own genetic information system. In general, the metazoan mitogenome is a closed, circular DNA molecule, ranging from 12 to 20 kb in length and usually containing 37 genes: 13 protein-coding genes (PCGs) (*atp6, atp8, cox1-3, cytb, nad1-6* and *nad4l*) of the respiratory chain, 2 ribosomal RNA (rRNA) genes (*rrnS* and *rrnL*) and 22 transfer RNA (tRNA) genes [2]. In addition, there are several noncoding regions in the mitogenome, and the longest noncoding “AT-rich” region is known as the control region (CR), which includes elements controlling the initiation and regulation of transcription and replication [3]. Owing to maternal inheritance, variable gene order, a low frequency of gene recombination and different genes having different evolutionary rates, mitochondrial sequences are widely used for species identification, genetic diversity assessment and phylogenetics at various taxonomic levels [4–7].

Since the discovery of cold seeps and hydrothermal vents in the deep sea, the unique biological communities that depend on chemosynthetic primary production have attracted the attention of researchers [8–11]. These deep-sea environments lack sunlight and exhibit high pressure, low oxygen and high levels of chemical toxicity due to various heavy metals, and the organisms that live there show a series of adaptations compared with marine species in coastal environments [12–15]. Mitochondria are the energy metabolism centers of eukaryotic cells, which can generate more than 95% of cellular energy through oxidative phosphorylation (OXPHOS) [3]. Therefore, mitochondrial PCGs may undergo evolutionary selection in response to metabolic requirements in extremely harsh environments. Numerous studies have found clear and compelling evidence of adaptive evolution in the mitogenome of organisms from extreme habitats, including Tibetan humans [16], Chinese snub-nosed monkeys [17], Tibetan horses [18–19], Tibetan wild yaks [20], galliform birds [21], and Tibetan loaches [22].

The family Vesicomyidae (Dall & Simpson, 1901) is widely distributed worldwide from shelf to hadal depths and comprises specialized bivalves occurring in reducing environments such as hydrothermal vents located in mid-ocean ridges and back-arc basins, cold seeps at continental margins and whale falls [23–26]. Studies have shown that vesicomyid bivalves rely upon the symbiotic chemoautotrophic bacteria in their gills for all or part of nutrition [27–28]. Based on the shells and soft body, the Vesicomyidae is divided into two subfamilies: Vesicomyinae and Pliocardiinae. The Vesicomyinae includes only one genus, *Vesicomya*, while Pliocardiinae currently contains 20 genera. Among the 20 genera, *Calyptogena* is the most diverse group of deep-sea vesicomyid bivalves in the western Pacific region and its marginal seas [29]. As some of the dominant species in the deep sea, vesicomyids are an interesting taxon with which to study the mechanisms of adaptation to diverse stressors in deep-sea habitats. Considering that the mitogenome has highly compact DNA and is easily accessible, several complete/nearly complete mitogenomes of vesicomyids have been sequenced [30–33] in recent years; however, limited information is available about the mechanism of adaptation to deep-sea habitats in vesicomyids at the mitogenome level.

In the present study, we obtained the mitogenome of *Calyptogena marissinica*, a new species of the family Vesicomyidae from the Haima cold seep of the South China Sea. First, the mitogenome organization, codon usage, and gene order information were obtained, and we compared the composition of this mitogenome with that of other available vesicomyid bivalve mitogenomes. Second, based on mitochondrial PCGs and 2 rRNA genes, the phylogenetic relationships between *C. marissinica* and other species from subclass Heterodonta were examined. Finally, to understand the adaptive evolution of deep-sea organisms, we conducted positive selection analysis of vesicomyid bivalve mitochondrial PCGs.

## Materials and Methods

### Sampling, identification and DNA extraction

Specimens of *C. marissinica* were sampled from the “Haima” methane seep in the northern sector of the South China Sea at a depth of 1,380-1,390 m using a remotely operated vehicle (ROV) in May 2018. Species-level morphological identification was performed according to the main points of Chen et al. (2018) [29]. Specimens were preserved at −80°C.until DNA extraction. Total genomic DNA was extracted using a DNeasy tissue kit (Qiagen, Beijing, China) following the manufacturer’s protocols.

### Illumina sequencing, mitogenome assembly and annotation

After DNA isolation, 1 μg of purified DNA was fragmented, used to construct short-insert libraries (insert size of 430 bp) according to the manufacturer’s instructions (Illumina), and then sequenced on an Illumina HiSeq 4000 instrument (San Diego, USA).

Prior to assembly, raw reads were filtered. This filtering step was performed to remove the reads with adaptors, the reads showing a quality score below 20 (Q<20), the reads containing a percentage of uncalled bases (“N” characters) equal to or greater than 10% and the duplicated sequences. The mitochondrial genome was reconstructed using a combination of *de novo* and reference-guided assemblies, and the following three steps were used to assemble the mitogenome. First, the filtered reads were assembled into contigs using SOAPdenovo 2.04 [34]. Second, contigs were aligned to the reference mitogenomes from species of the family Vesicomyidae using BLAST, and aligned contigs (≥80% similarity and query coverage) were ordered according to the reference mitogenomes. Third, clean reads were mapped to the assembled draft mitogenome to correct the incorrect bases, and the majority of gaps were filled via local assembly.

The mitochondrial genes were annotated using homology alignments and *de novo* prediction, and EVidenceModeler [35] was used to integrate the gene set. rRNA genes and tRNA genes were predicted by rRNAmmer [36] and tRNAscan-SE [37]. A whole-mitochondrial genome BLAST search (E-value ≤ 1e^-5^, minimal alignment length percentage ≥ 40%) was performed against 5 databases, namely, the KEGG (Kyoto Encyclopedia of Genes and Genomes), COG (Clusters of Orthologous Groups), NR (Non-Redundant Protein), Swiss-Prot and GO (Gene Ontology) databases. Organellar Genome DRAW [38] was used for circular display of the *C. marissinica* mitogenome.

### Sequence analysis

The nucleotide composition and codon usage were computed using DnaSP 5.1 [39]. The AT and GC skews were calculated with the following formulas: AT skew = (A − T) / (A + T) and GC skew = (G − C) / (G + C) [40], where A, T, G and C are the occurrences of the four nucleotides. Tandem Repeats Finder 4.0 [41] was used to search for the tandem repeat sequences. The online version of Mfold [42] was applied to infer potential secondary structure, and if more than one secondary structure appeared, the one with the lowest free energy score was used.

### Phylogenetic analysis

The phylogeny of the subclass Heterodonta was reconstructed using mitogenome data from 41 species, including 2 Lucinida species, 2 Myoida species, and 37 Veneroida species, and *Chlamys farreri* and *Mimachlamys nobilis* from the subclass Pteriomorphia served as outgroups (S1 Table). Our data set was based on nucleotide and amino acid sequences from 9 mitochondrial PCGs (*cox1, cox2, cox3, cob, atp6, nad1, nad4, nad5*, and *nad6*) and 2 rRNA genes. The *atp8, nad2, nad4l* and *nad6* sequences were excluded due to several species with incomplete mitogenomes. Multiple alignments of the 11 genes were conducted using MUSCLE 3.8.31, followed by manual correction. In aligned sequences, ambiguously aligned regions and gaps were removed using Gblocks server ver. 0.91b [43] with the default setting. ModelTest 2.1.10 [44] and ProtTest 3.4 [45] were used to select the best-fit evolutionary models GTR + I + G and LG + I + G + F for the nucleotide dataset and amino acid dataset, respectively. Maximum likelihood (ML) analysis was performed using RAxML ver. 8.2.8 [46]. Topological robustness for the ML analysis was evaluated with 100 replicates of bootstrap support. Bayesian inference (BI) was conducted using MrBayes 3.2.6 [47], and four Markov chain Monte Carlo (MCMC) chains were run for 10^6^ generations, with sampling every 100 generations and a 25% relative burn-in. All phylogenetic trees were graphically edited with the iTOL 3.4.3 (https://itol.embl.de/itol.cgi).

### Positive selection analysis

Comparing the nonsynonymous/synonymous substitution ratios (ω = *dN*/*dS*) has become a useful approach for quantifying the impact of natural selection on molecular evolution [48]. ω >1, =1 and <1 indicate positive selection, neutrality and purifying selection, respectively. The codon-based maximum likelihood (CodeML) method implemented in the PAML package [49] was applied to estimate the *dN*/*dS* ratio ω. The combined database of 13 mitochondrial PCGs was used for the selection pressure analyses. Both the ML and Bayesian phylogenetic trees were separately used as the working topology in all CodeML analyses.

To evaluate positive selection in the vesicomyid bivalves, we used branch models in the present study. First, a one-ratio model (M_0_), the simplest model, which allows only a single ω ratio for all branches in the phylogeny [50], was used to preliminarily estimate the ω value for the gene sequences. Then, a two-ratio model, which allows the background lineages and foreground lineages to have different ω ratio values, was used to detect positive selection acting on branches of interest [51–52]. Last, a free-ratio model, which allows ω ratio variation among branches, was used to estimate ω values on each branch [52]. Here, a one-ratio model and a free-ratio model were compared to confirm whether different clades in Heterodonta had different ω values. Additionally, we compared a one-ratio model and a two-ratio model to investigate whether deep-sea vesicomyid clades are subjected to more selection pressure than other Heterodonta species in coastal waters. ω_0_ and ω_1_ represent the values for the other Heterodonta clades in the phylogeny and the vesicomyid clades, respectively. Pairwise models were compared with critical values of the Chi square (χ^2^) distribution using likelihood ratio tests (LRTs), in which the test statistic was estimated as twice the log likelihood (2ΔL) and the degrees of freedom were estimated as the difference in the number of parameters for each model.

Furthermore, we fit branch-site models to examine positive selection on some sites among the vesicomyid clades. Branch-site models allow ω to vary both among sites in the protein and across branches on the tree. Branch-site model A (positive selection model) was used to identify the positively selected sites among the lineages of vesicomyids (marked as foreground lineages). The presence of sites with ω > 1 suggests that model A fits the data significantly better than the corresponding null model. Bayes Empirical Bayes analysis was used to calculate posterior probabilities in order to identify sites under positive selection on the foreground lineages if the LRTs was significant [53].

## Results and Discussion

### *C. marissinica* mitogenome organization

The Illumina HiSeq runs resulted in 20,359,890 paired-end reads from the *C. marissinica* library with an insert size of approximately 450 bp. Selective-assembly analysis showed that 2,422 Mb of clean data (Q20 quality score of 97.01%) was assembled into a 17,374-bp circular molecule, which represented the complete mitogenome of *C. marissinica* (Fig 1 and Table 1). This length is shorter than that of the complete mitogenome of other vesicomyid bivalves, which ranges from 19,738 bp in *Calyptogena magnifica* [30] to 19,424 bp in *Abyssogena phaseoliformis* [32]. The genome encodes 37 genes, including 13 PCGs, 2 rRNA genes, and 22 tRNA genes (duplication of *tRNA*^*Leu*^ and *tRNA*^*Ser*^). All of the genes are encoded on the heavy (H) strand, as consistently reported for other bivalves [32–33,54], and transcribed in the same direction. A total of 2,287 bp of noncoding nucleotides are scattered among 23 intergenic regions varying from 1 to 1,676 bp in length (Table 1). The largest noncoding region (1,676 bp) is located between *tRNA*^*Trp*^ and *nad6* and is identified as the putative control region (CR) due to its location and high A+T content (73.3%). Furthermore, there are four overlaps between adjacent genes in the *C. marissinica* mitogenome with a size range of 1 to 5 bp (*tRNA*^*Glu*^ / *tRNA*^*Ser*(*UCA*)^, *tRNA*^*Leu*(*UUA*)^ / *nad1, rrnS* / *tRNA*^*Met*^, and *cox3* / *tRNA*^*Phe*^).

**Fig 1.**
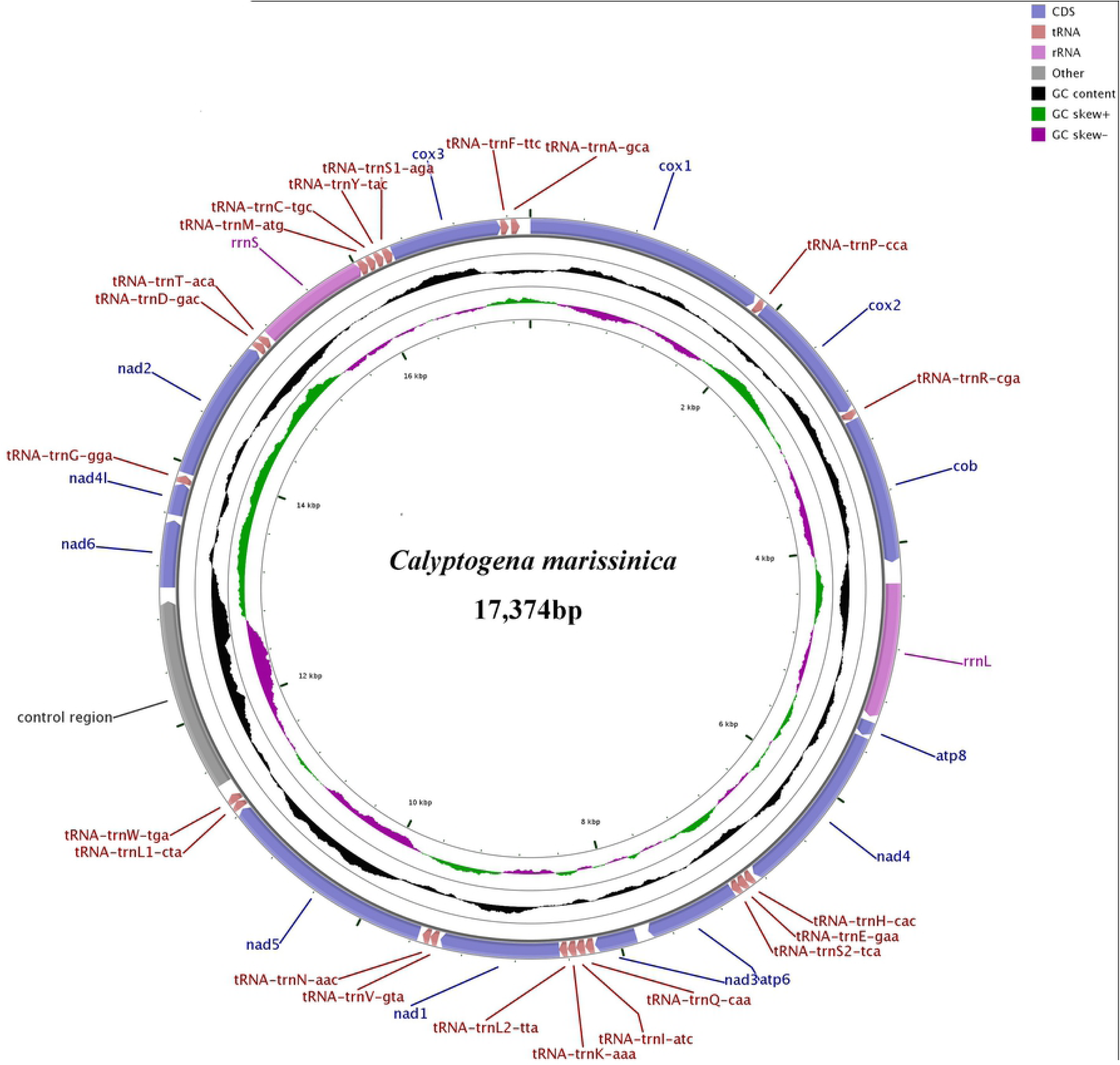
Complete mitogenome map of *C. marissinica*. All 37 genes are encoded on the heavy (H) strand. Genes for proteins and rRNAs are shown with standard abbreviations. Genes for tRNAs are displayed by a single letter for the corresponding amino acid, with two leucine tRNAs and two serine tRNAs differentiated by numerals.

**Table 1.** Mitogenome organization of *C. marissinica*.

| Name | Strand | Range | Size |  | Codon |  |  | Intergenic nucleotides |
| --- | --- | --- | --- | --- | --- | --- | --- | --- |
|  |  |  | Nucleotides | Amino acids | Start | Stop | Anticodon |  |
| *cox1* | + | 1-1833 | 1833 | 610 | ATG | TAA |  | - |
| *tRNA-Pro* | + | 1854-1917 | 64 |  |  |  | TGG | 20 |
| *cox2* | + | 1918-2934 | 1017 | 338 | ATG | TAA |  | 0 |
| *tRNA-Arg* | + | 2941-3005 | 65 |  |  |  | TCG | 6 |
| *cob* | + | 3010-4143 | 1134 | 377 | ATG | TAA |  | 4 |
| *rrnL* | + | 4305-5340 | 1036 |  |  |  |  | 161 |
| *atp8* | + | 5377-5493 | 117 | 38 | --- | TAG |  | 36 |
| *nad4* | + | 5506-6852 | 1347 | 448 | ATG | TAA |  | 12 |
| *tRNA-His* | + | 6873-6933 | 61 |  |  |  | GTG | 20 |
| *tRNA-Glu* | + | 6934-6999 | 66 |  |  |  | TTC | 0 |
| *tRNA-Ser<sup>UCA</sup>* | + | 6996-7059 | 64 |  |  |  | TGA | -4 |
| *atp6* | + | 7060-7773 | 714 | 237 | ATG | TAA |  | 0 |
| *nad3* | + | 7872-8192 | 321 | 106 | ATT | TAA |  | 98 |
| *tRNA-Gln* | + | 8203-8269 | 67 |  |  |  | TTG | 10 |
| *tRNA-Ile* | + | 8272-8338 | 67 |  |  |  | GAT | 2 |
| *tRNA-Lys* | + | 8339-8405 | 67 |  |  |  | TTT | 0 |
| *tRNA-Leu<sup>UUA</sup>* | + | 8407-8469 | 63 |  |  |  | TAA | 1 |
| *nad1* | + | 8467-9381 | 915 | 304 | ATA | TAG |  | -3 |
| *tRNA-Val* | + | 9399-9460 | 62 |  |  |  | TAC | 17 |
| *tRNA-Asn* | + | 9461-9522 | 62 |  |  |  | GTT | 0 |
| *nad5* | + | 9550-11223 | 1674 | 557 | ATA | TAA |  | 27 |
| *tRNA-Leu<sup>CUA</sup>* | + | 11236-11297 | 62 |  |  |  | TAG | 12 |
| *tRNA-Trp* | + | 11298-11362 | 65 |  |  |  | TCA | 0 |
| *control region* | + | 11363-13038 | 1676 |  |  |  |  | 0 |
| *nad6* | + | 13039-13554 | 516 | 171 | ATT | TAA |  | 0 |
| *nad4l* | + | 13595-13843 | 249 | 82 | ATT | TAA |  | 40 |
| *tRNA-Gly* | + | 13844-13907 | 64 |  |  |  | TCC | 0 |
| *nad2* | + | 13925-15010 | 1086 | 361 | ATT | TAG |  | 17 |
| *tRNA-Asp* | + | 15020-15081 | 62 |  |  |  | GTC | 9 |
| *tRNA-Thr* | + | 15082-15142 | 61 |  |  |  | TGT | 0 |
| *rrnS* | + | 15160-16027 | 868 |  |  |  |  | 17 |
| *tRNA-Met* | + | 16023-16089 | 67 |  |  |  | CAT | -5 |
| *tRNA-Cys* | + | 16092-16153 | 62 |  |  |  | GCA | 2 |
| *tRNA-Tyr* | + | 16159-16220 | 62 |  |  |  | GTA | 5 |
| *tRNA-Ser<sup>AGA</sup>* | + | 16228-16296 | 69 |  |  |  | TCT | 7 |
| *cox3* | + | 16297-17148 | 852 | 283 | ATG | TAG |  | 0 |
| *tRNA-Phe* | + | 17148-17210 | 63 |  |  |  | GAA | -1 |
| *tRNA-Ala* | + | 17228-17293 | 66 |  |  |  | TGC | 17 |

The *C. marissinica* mitogenome has a nucleotide composition of 25.9% A, 10.8% C, 23.8% G, and 39.5% T and an overall AT content of 65.4%. The AT skew and GC skew are well conserved among vesicomyids, which vary from −0.165 to −0.230 and 0.343 to 0.440, respectively (Table 2). For the *C. marissinica* mitogenome, the AT skew is −0.209, and the GC skew is 0.375, which indicates bias toward T and G similar to that in other vesicomyids. The complete mitochondrial DNA sequence has been deposited in GenBank under accession number MK948426.

**Table 2.** Mitogenomes of Vesicomyidae species sequenced to date and their genomic features.

| Species | Genus | Accession number | Length (bp) | Genome |  |  | Protein-coding genes |  |  | rrnL |  | rrnS |  | tRNAs |  | Control region |  |
| --- | --- | --- | --- | --- | --- | --- | --- | --- | --- | --- | --- | --- | --- | --- | --- | --- | --- |
|  |  |  |  | AT% | AT skew | GC skew | Length (aa) | AT% (all) | AT% (3rd) | Length (bp) | AT% | Length (bp) | AT% | Number/Length(bp) | AT% | Length (bp) | AT% |
| *Abyssogena mariana*<sup>1</sup> | *Abyssogena* | LC126311 | 15,927* | 69.8 | -0.210 | 0.408 | 3884 | 69.0 | 73.9 | 1196 | 71.2 | 862 | 70.8 | 23/1279 | 71.7 | - | - |
| *Abyssogena phaseoliformis* | *Abyssogena* | AP014557 | 19,424 | 70.4 | -0.199 | 0.440 | 3881 | 68.3 | 73.7 | 1196 | 71.2 | 862 | 70.9 | 24/1282 | 70.4 | 3438 | 74.4 |
| *Akebiconcha kawamurai*<sup>2</sup> | *Akebiconcha* | AP014551 | 12,946* | 65.2 | -0.222 | 0.371 | 3249 | 62.5 | 68.7 | 1194 | 69.7 | 205 | 67.7 | 17/1090 | 69.0 | - | - |
| *Archivesica gigas*<sup>1</sup> | *Archivesica* | MF959623 | 15,674* | 65.0 | -0.228 | 0.389 | 3878 | 63.6 | 70.6 | 1223 | 68.9 | 879 | 67.9 | 21/1212 | 68.7 | - | - |
| *Archivesica pacifica*<sup>1</sup> | *Archivesica* | MF959624 | 17,782* | 68.6 | -0.214 | 0.429 | 4002 | 67.1 | 79.5 | 1226 | 71.3 | 885 | 69.9 | 22/1282 | 69.6 | - | - |
| *Archivesica* sp.<sup>1</sup> | *Archivesica* | MF959622 | 15,650* | 64.8 | -0.228 | 0.386 | 3889 | 63.7 | 70.8 | 1221 | 69.0 | 879 | 67.8 | 20/1214 | 68.7 | - | - |
| *Calypptogena fausta*<sup>2</sup> | *Calypptogena* | AP014549 | 13,509* | 66.0 | -0.218 | 0.394 | 3410 | 64.7 | 70.7 | 1189 | 70.3 | 205 | 67.8 | 17/1092 | 70.0 | - | - |
| *Calypptogena laubirei*<sup>2</sup> | *Calypptogena* | AP014553 | 12,968* | 64.3 | -0.226 | 0.361 | 3259 | 61.6 | 67.1 | 1191 | 69.0 | 204 | 67.2 | 17/1090 | 67.8 | - | - |
| *Calypptogena magnifica* | *Calypptogena* | NC_028724 | 19,738 | 68.4 | -0.195 | 0.390 | 3928 | 65.5 | 75.6 | 1219 | 70.5 | 935 | 70.0 | 22/1347 | 70.2 | 3910 | 75.2 |
| *Calypptogena marissinica* | *Calypptogena* |  | 17,374 | 65.4 | -0.209 | 0.375 | 3912 | 63.2 | 69.7 | 1,036 | 67.2 | 868 | 67.7 | 22/1,411 | 68.7 | 1676 | 73.3 |
| *Calypptogena nautilei*<sup>2</sup> | *Calypptogena* | AP014554 | 13,281* | 69.4 | -0.196 | 0.353 | 3298 | 68.0 | 74.4 | 1182 | 72.1 | 204 | 72.1 | 17/1088 | 71.4 | - | - |
| *Calypptogena pacifica*<sup>2</sup> | *Calypptogena* | AP014556 | 13,454* | 67.6 | -0.222 | 0.420 | 3390 | 66.6 | 72.6 | 1195 | 70.8 | 204 | 69.6 | 17/1088 | 69.5 | - | - |
| *Isorropodon fossajaponicum*<sup>1</sup> | *Isorropodon* | AP014550 | 19,556* | 68.2 | -0.165 | 0.343 | 3894 | 66.6 | 70.6 | 1199 | 70.3 | 861 | 68.3 | 24/1290 | 70.3 | - | - |
| *Phreagena kilmeri*<sup>2</sup> | *Phreagena* | AP014552 | 12,944* | 64.9 | -0.223 | 0.365 | 3249 | 63.4 | 68.0 | 1191 | 69.5 | 204 | 68.3 | 17/1089 | 68.7 | - | - |
| *Phreagena okutanii*<sup>1</sup> | *Phreagena* | AP014555 | 16,336* | 65.6 | -0.230 | 0.405 | 3833 | 64.0 | 68.1 | 1191 | 69.7 | 861 | 67.7 | 23/1277 | 68.9 | - | - |
| *Phreagena soyoe*<sup>2</sup> | *Phreagena* | AP014558 | 12,941* | 64.9 | -0.223 | 0.365 | 3249 | 63.4 | 67.9 | 1190 | 69.5 | 204 | 68.1 | 17/1089 | 68.7 | - | - |
| *Pliocardia stearnsii*<sup>2</sup> | *Pliocardia* | AP014559 | 13,012* | 67.7 | -0.230 | 0.402 | 3265 | 66.6 | 72.7 | 1201 | 71.6 | 203 | 69.5 | 17/1096 | 69.1 | - | - |

Note: \* indicates incomplete mitogenomes.
<sup>1</sup> Incomplete mitogenomes for which the control region was not sequenced.
<sup>2</sup> Incomplete mitogenomes for which *nad2*, *nad4l*, *nad6*, a few tRNA genes and the control region were not sequenced.

### Protein-coding genes

The total length of all 13 PCGs of *C. marissinica* is 11,775 bp, accounting for 67.8% of the complete length of the mitogenome, and the PCGs encode 3,912 amino acids (Table 2). In the mitogenome of metazoans, most PCGs start with the standard ATN codon [2,55–56]. In the present study, with the exception of the *atp8* gene, which had the alternate initiation codon GTG, all the PCGs were initiated by typical ATN codons: 6 genes (*atp6, cob, cox1, cox2, cox3*, and *nad4*) were initiated by ATG, 4 genes (*nad2, nad3, nad4l*, and *nad6*) were initiated by ATT, and 2 genes (*nad1* and *nad5*) were initiated by ATA. Notably, genes are commonly initiated by GTG in vesicomyid bivalves [31], and the amino acid encoded by GTG is valine, which belongs to the nonpolar amino acids, such as methionine and isoleucine encoded by ATN. Moreover, in eight other vesicomyid bivalves (*Archivesica* sp., *Archivesica gigas, Archivesica pacifica, C. magnifica, Abyssogena mariana, Ab. phaseoliformis, Isorropodon fossajaponicum*, and *Phreagena okutanii*), *cox3* had a truncated termination codon, TA [31]. Previous studies have shown that the truncated stop codon is common in the metazoan mitogenome and might be corrected by posttranscriptional polyadenylation [57–58]. However, in the mitogenome of *C. marissinica*, all of the PCGs were ended by a complete TAA (*atp6, cob, cox1, cox2, nad3, nad4, nad4l, nad5*, and *nad6*) or TAG (*atp8, cox3, nad1*, and *nad2*) termination codon.

Numerous studies have indicated that metazoan mitogenomes usually have a bias toward a higher representation of nucleotides A and T, leading to a subsequent bias in the corresponding encoded amino acids [56,59–61]. In the mitogenome of *C. marissinica*, the A+T contents of PCGs and third codon positions are 63.2% and 69.7%, respectively, which are similar to the values observed in other vesicomyids (Table 2). The amino acid usage and relative synonymous codon usage (RSCU) values in the PCGs of *C. marissinica* are summarized in Fig 2. The mitogenome encodes a total of 3,912 amino acids, among which leucine (13.6%) and glutamine (1.4%) are the most and the least frequently used, respectively. As mentioned earlier, the amino acids encoded by A+T-rich codon families (Asn, Ile, Lys, Met, Phe and Tyr) have a higher frequency of use than those encoded by G+C-rich codon families (Ala, Arg, Gly and Pro). The RSCU values indicate that the six most commonly used codons are TTA (Leu), ACT (Thr), GGG (Gly), TCT (Ser), GCT (Ala), and CCT (Pro) (Fig 2), which show A+T bias at their third codon position. In addition, the codons with A and T in the third position are used more frequently than other synonymous codons. These features reflect codon usage with A and T biases at the third codon position, which are similar to the biases that exist in many metazoans [62–65].

**Fig 2.**
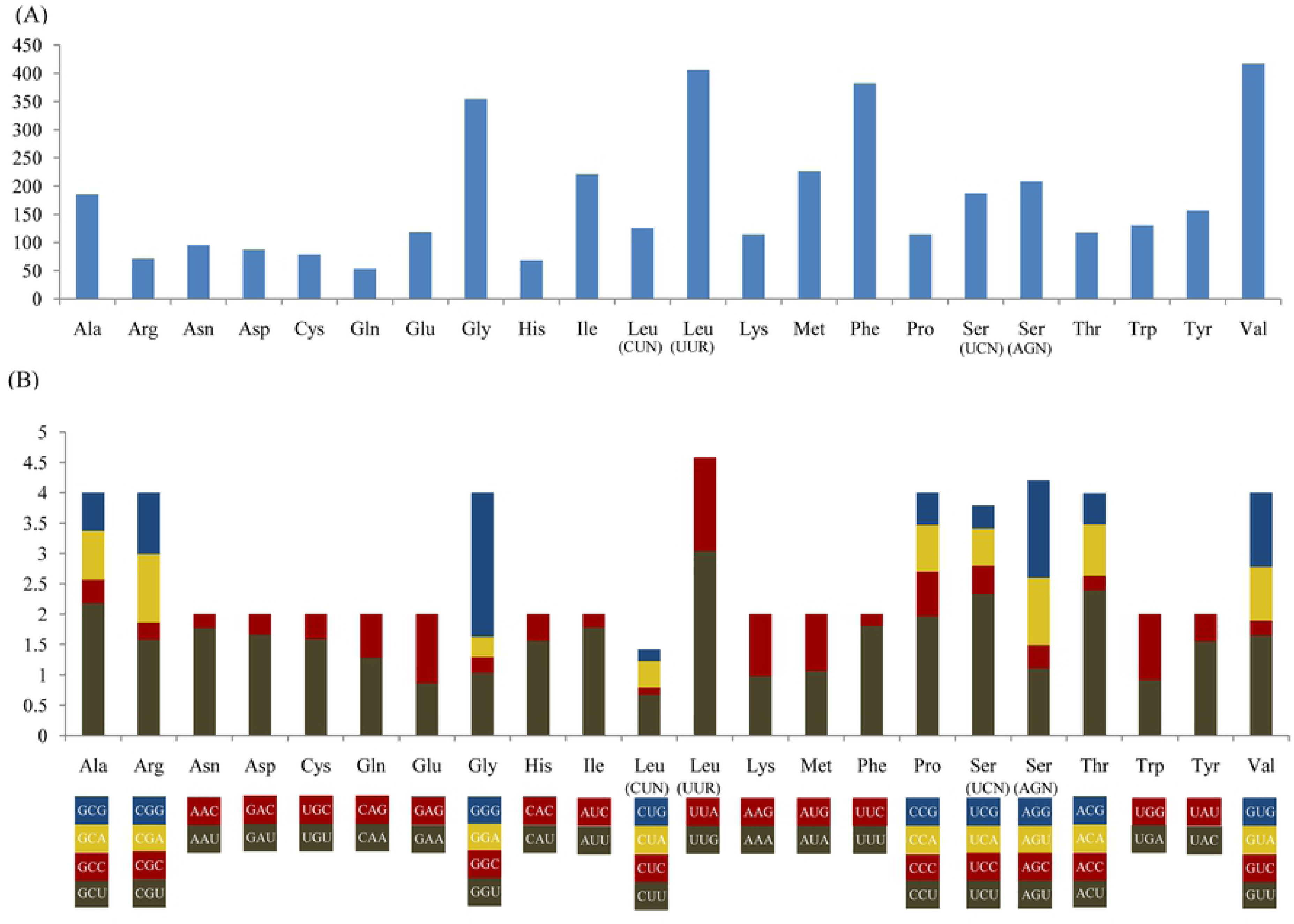
Codon usage (A) and the relative synonymous codon usage (RSCU) (B) of the *C. marissinica* mitogenome. Numbers to the left refer to the total number of codons (A) and the RSCU values (B). Codon families are provided on the X axis.

### Ribosomal RNA and transfer RNA genes

The *rrnL* and *rrnS* genes of *C. marissinica* are 1,036 bp (AT% = 67.2) and 868 bp (AT% = 67.7) in length, respectively. As in other vesicomyid bivalves, *rrnL* is located between the *cytb* and *atp8* genes, and *rrnS* is located between *tRNA*^*Thr*^ and *tRNA*^*Met*^. The largest known *rrnL* and *rrnS* genes are 1,226 bp in *Ar. pacific* and 935 bp in *C. magnifica*, respectively [30–32].

Twenty-two tRNA genes were identified in the mitogenome of *C. marissinica*, which is typical for metazoans. However, the number of tRNA genes varies among other vesicomyid bivalves (Table 2). The length of tRNA genes in *C. marissinica* ranges from 61 (*tRNA*^*His*^ and *tRNA*^*Thr*^) to 69 (*tRNA*^*Ser (AGA)*^) bp (Table 1), and the AT content of the tRNA genes is 68.7%. The secondary structures of tRNA genes are schematized in S1 Fig. Generally, a typical tRNA clover-leaf structure includes a 7-8 bp aminoacyl acceptor stem, a 3-5 bp TψC stem, a 5 bp anticodon stem and a 4 bp DHU stem. In the present study, most of the tRNA genes had the typical secondary structure, except for *tRNA*^*His*^, *tRNA*^*Thr*^, *tRNA*^*Tyr*^, *tRNA*^*Ser(UCA)*^ and *tRNA*^*Ser(AGA)*^. In *tRNA*^*His*^, *tRNA*^*Thr*^and *tRNA*^*Tyr*^, the TψC loops are incomplete, which is not observed in other vesicomyid bivalves [31–33], and this feature might be a specific character in the *C. marissinica* mitogenome. In *tRNA*^*Ser(UCA)*^ and *tRNA*^*Ser(AGA)*^, the DHU stems are reduced to a simple loop, as in many other bivalve mitogenomes [31,66]. Many studies have shown that the incomplete clover-leaf secondary structure of tRNA genes is common in metazoan mitogenomes and that aberrant tRNA genes can still function normally through posttranscriptional RNA editing and/or coevolved interacting factors [67–69]. Additionally, several mismatch pairs were detected within amino acid acceptors and anticodon stems in tRNA genes of *C. marissinica*. Such mismatches seem to be ubiquitous phenomena in the mitogenomes of many organisms and can also be corrected by posttranscriptional RNA editing [56,64,70–71].

### Noncoding regions and gene arrangement

A total of 23 noncoding regions (totaling 2,216 bp) are distributed in the *C. marissinica* mitogenome. The longest noncoding region (1,676 bp), located between *tRNA*^*Trp*^ and *nad6*, corresponds to the control region identified in most other vesicomyids. The nucleotide content of the 1,676 bp control region is 34.25% A, 39.02% T, 16.29% G, and 10.44% C. The A + T content (73.27%) of this region is higher than that of other regions in the *C. marissinica* mitogenome (Table 2). In general, the mitochondrial control region is subjected to less evolutionary pressure than PCGs and thus has the highest variation in the whole mitogenome [72–73].

Additionally, in the mitochondrial control region of *C. marissinica*, we found a tandemly arranged repeated sequence, which was 354 bp in length (positions 12,675-13,028), including three identical tandem repeat units of 118 bp (Fig 3). The tandem repeat sequence could be folded into stem-loop secondary structures with minimized free energy (Fig 3), which is a common phenomenon in invertebrates [61,64–65,74]. The control region in the mitogenome is essential for transcription and replication in animals [75–76]. Therefore, the stem-loop structures mentioned above may play an important role in gene replication and regulation. In addition, some other peculiar patterns, such as special “G(A)_n_T” motifs and AT-rich sequences, were observed in the control region of the *C. marissinica* mitogenome (Fig 3). Furthermore, similar characteristics (e.g., repetitive elements, G(A)_n_T motifs and AT-rich sequences) were also observed in the deep-sea anemone *Bolocera* sp., alvinocaridid shrimp *Shinkaicaris leurokolos* and spongicolid shrimp *Spongiocaris panglao* [61,64–65]. In view of the particularity of the deep-sea environment, we speculate that these special control region elements are involved in the regulation of replication and transcription of the mitogenome and help organisms adapt to extreme deep-sea habitats.

**Fig 3.**
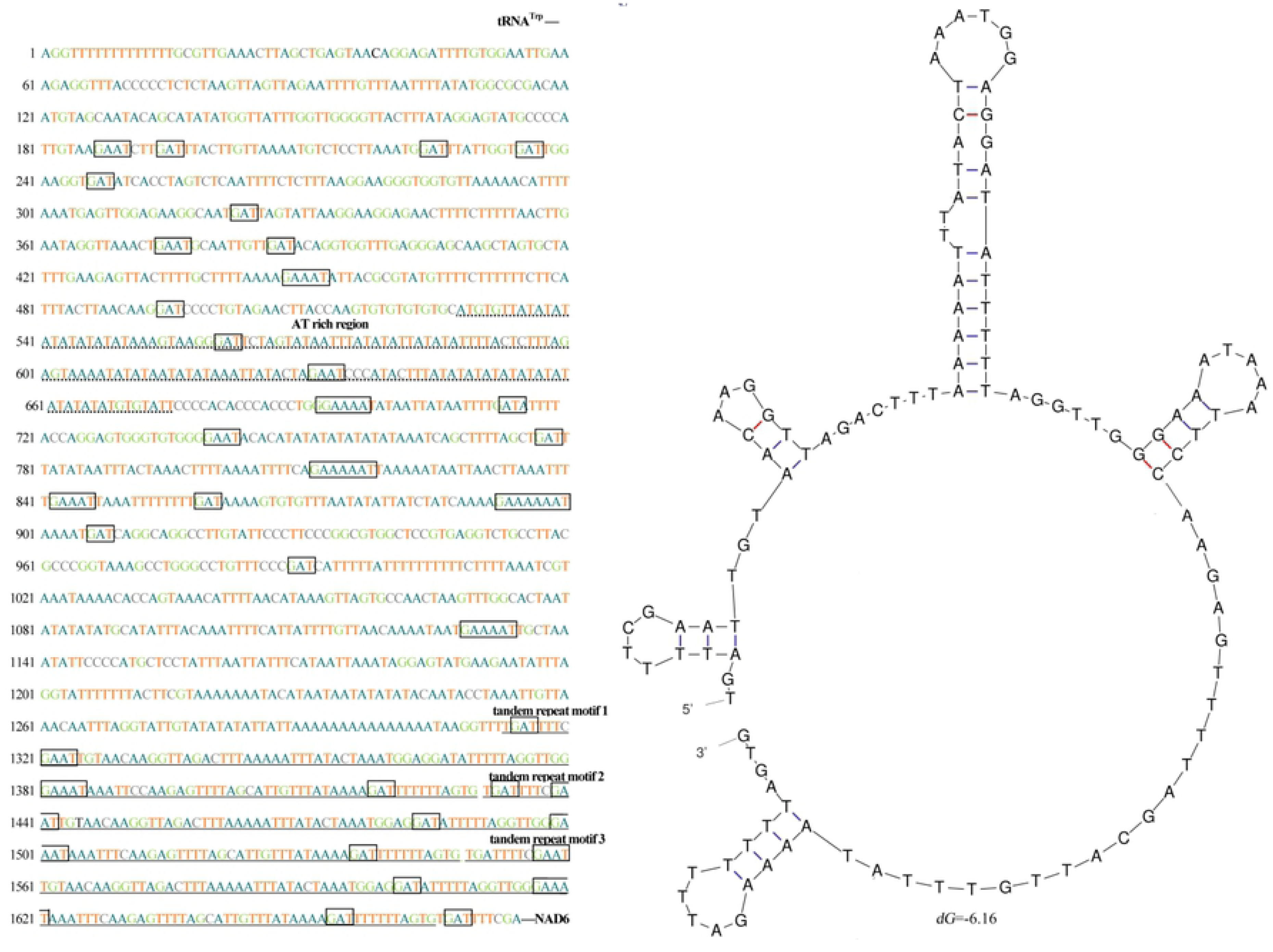
Nucleotide sequences and stem-loop structures of the tandem repeat motifs in the control region (CR) of the *C. marissinica* mitogenome. The CR is flanked by sequences encoding *tRNATrp* and *nad6*. The CR consists of certain patterns, such as special G(A)_n_T motifs (marked with a box), AT-rich regions and tandem repeat motifs.

In contrast to other metazoans, the Mollusca showed frequent and extensive variation in gene arrangement, and among them, bivalves showed more gene order variation in their mitogenomes [77–79]. Here, a comparison of the *C. marissinica* mitogenome with the other twelve Heterodonta mitogenomes is shown in Fig 4. All thirteen Heterodonta mitogenomes come from two orders (five families): Myoida (family Hiatellidae) and Veneroida (family Tellinidae, family Mactridae, family Veneridae and family Vesicomyidae). Among the Heterodonta mitogenomes analyzed in the present study, the gene arrangement has a distinct difference between the family Vesicomyidae and other species (Fig 4). In the family Vesicomyidae, we found that if the tRNA genes are not considered, the nine vesicomyid bivalves have a completely identical gene arrangement of PCGs. When compared to the “standard” mitogenome of *Ar. pacific, C. magnifica* and *C. marissinica*, several additional tRNA genes were identified in *Ab. mariana* (*tRNA*^*Leu3*^), *Ab. phaseoliformis* (*tRNA*^*His2*^ and *tRNA*^*Ser3*^), *I. fossajaponicum* (*tRNA*^*Asn2*^ and *tRNA*^*Lys2*^) and *Ph. okutanii* (*tRNA*^*Met2*^) (Fig 4). As a general rule, additional gene copies usually obtained by gene replication and different gene copies would share some sequence identity with each other. However, analysis showed that the aforementioned additional tRNA genes have low similarity to other tRNA genes that encode the same tRNAs [31]. The remolding of tRNA genes, DNA shuffling and the point mutations in the anticodons may all provide chances for tRNA gene rearrangement within mitogenomes [3,80–81]. Furthermore, gene rearrangements usually occurred around the control regions, which are considered the replication origins. Perhaps gene replication events occur frequently in this region, and consequently, more novel gene arrangements will be found in this region. To date, there are four known mechanisms of gene rearrangements in mitogenomes: inversion, transposition, reverse transposition and tandem duplication-random losses (TDRLs) [82–83]. However, the specific mechanism of significant differences in mitochondrial gene arrangements in mollusks has not been completely clarified. With the determination of mitogenomes in more species of this phylum, the mechanism of large-scale rearrangement of mitochondrial genes in mollusks will be identified by further comparing and summarizing the rules of gene arrangement among different species.

**Fig 4.**
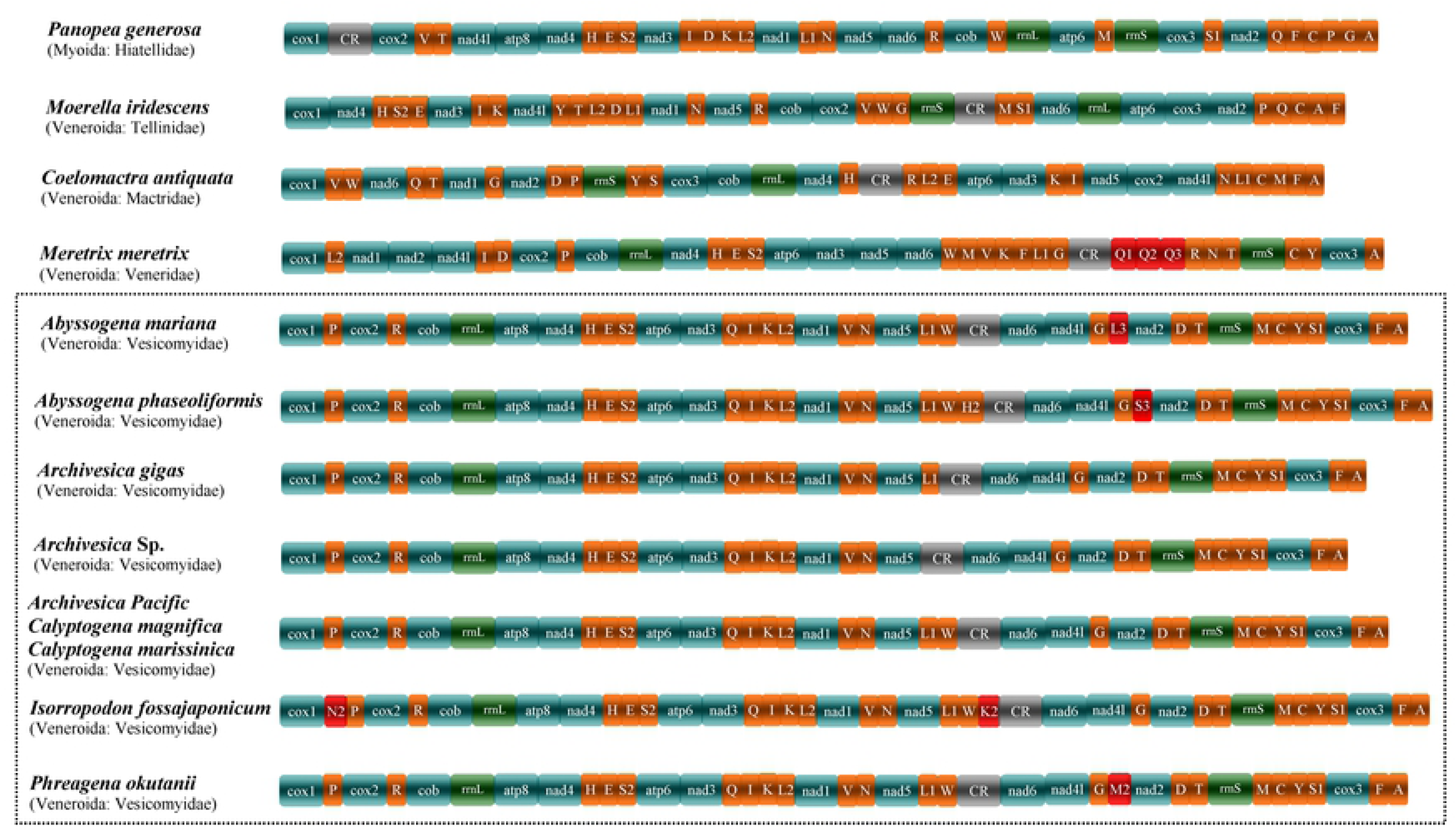
Mitochondrial gene arrangement of 13 species in the subclass Heterodonta (*Panopea generosa, Moerella iridescens, Coelomactra antiquata, Meretrix meretrix* and 9 vesicomyid clams). CR indicates the control region. Genes for tRNAs are displayed by a single letter for the corresponding amino acid, with two leucine tRNAs and two serine tRNAs differentiated by numerals. Uniquely derived gene positions of individual species are depicted in red. Sequence segments are not drawn to scale.

### Phylogenetic relationships

Since several vesicomyid bivalves have incomplete mitogenomes at present, phylogenetic analyses were performed based on nucleotide and amino acid sequences of 9 mitochondrial PCGs (*atp6, cox1, cox2, cox3, cob, nad1, nad3, nad4*, and *nad5*) and 2 rRNA genes by maximum likelihood (ML) and Bayesian inference (BI) methods (Fig 5, S1-S3 Fig). The tree topologies resulting from the BI and ML analyses were not the same. There are two potential reasons for this discrepancy: one is that the presence of noncoding rRNA genes made the databases of nucleotides and amino acids different, and the other is the fact that several clades are represented by only one or two species each. The phylogenetic analyses clustered *C. marissinica* with the previously reported vesicomyid bivalves with high support values (Fig 5). In all phylogenetic trees, the family Vesicomyidae first clustered well with Veneridae and then united with Mactridae, which corroborates earlier studies of phylogenetic relationships based on the concatenated 12 PCGs and 2 rRNA genes [31–33]. *Calyptogena* (*sensu lato*) is the most diverse group of deep-sea vesicomyid bivalves in the western Pacific region and its marginal seas. Until now, the composition, evolutionary position and level of the genus *Calyptogena* have been the subject of discussion [84–86]. Phylogenetic reconstruction using the cytochrome oxidase *c* subunit I (*cox1*) gene showed that *C. marissinica* was clearly nested within a fully supported monophyletic clade corresponding to *Calyptogena sensu lato* and consisting of all included *Calyptogena* (*sensu lato*) species [29]. Notably, in our studies, *C. marissinica* showed a close genetic relationship with the *Akebiconcha* species (Fig 5). Therefore, additional mitogenomes of a greater number of vesicomyid bivalves, combined with morphological characters, are necessary to determine the phylogenetic relationships among members of this family.

**Fig 5.**
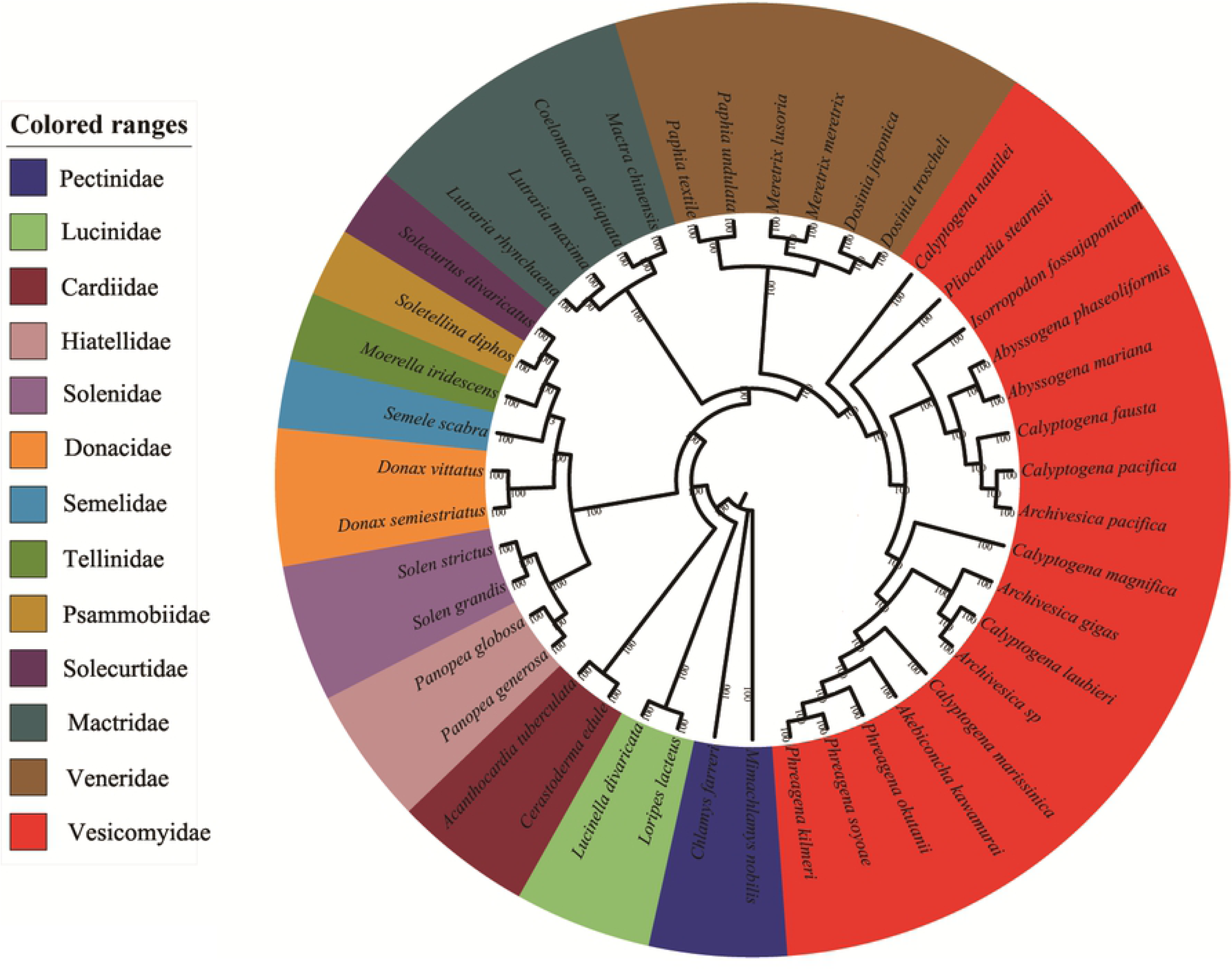
Phylogenetic tree derived from Bayesian analyses based on concatenated nucleotide sequences of 9 mitochondrial PCGs (*cox1, cox2, cox3, cob, atp6, nad1, nad4, nad5*, and *nad6*) and 2 ribosomal RNA genes (*rrnS* and *rrnL*). Numbers on branches are Bayesian posterior probabilities (percent). Two Pectinidae species belonging to the subclass Pteriomorphia were used as outgroups.

### Positive selection analysis

Purifying selection is the predominant force in the evolution of mitogenomes, but because mitochondria are the main sites of aerobic respiration and are essential for energy metabolism, weak and/or episodic positive selection may occur against this background of strong purifying selection under reduced oxygen availability or greater energy requirements [87–88]. As proven by many studies, mitochondrial PCGs underwent positive selection in animals that survived in hypoxic environments or had higher energy demands for locomotion, such as Tibetan humans, Ordovician bivalves, diving cetaceans and flying insects [16,89–91].

Considering that the special habitats of the deep sea may impact the function of mitochondrial genes, we examined potential positive selection in the Vesicomyidae lineage using CodeML from the PAML package. Although different tree-building methods were used, the results of positive selection analyses were generally consistent (Table 3). In the analysis of branch models, the ω (*dN/dS*) ratio calculated under the one-ratio model (M_0_) was 0.02272 for the 13 mitochondrial PCGs of sampled Heterodonta bivalves, suggesting that these genes have experienced constrained selection pressure to maintain function. Then, in the comparison of the one-ratio model (M_0_) and the free-ratio model, the LRTs indicated that the free-ratio model fit the data better than the one-ratio model (Table 3), which means that the ω ratios are indeed different among lineages. Furthermore, the two-ratio model was also found to fit the data better than the one-ratio model (Table 3) when the family Vesicomyidae was set as a foreground branch. The ω ratio of the Vesicomyidae branch was almost treble that of other branches (ω_1_ = 0.06398 and ω_0_ = 0.02278), indicating divergence in selection pressure between vesicomyid bivalves and other shallow-sea Heterodonta species. However, the ω ratio of the family Vesicomyidae (ω_1_= 0.06398) was still significantly less than 1. This result is consistent with the known functional significance of mitochondria as a respiration chain necessary for electron transport and OXPHOS [92].

**Table 3.** CODEML analyses of selection pressure on mitochondrial genes in the Vesicomyidae lineage.

| Trees | Models | lnL | Parameter estimates | Models compared | 2ΔL | *p*-value |
| --- | --- | --- | --- | --- | --- | --- |
| Branch models |  |  |  |  |  |  |
| Bayesian tree | M0 | -203672.1001 | $\omega = 0.02272$ | | | |
| | Two-ratio | -203666.6298 | $\omega_0 = 0.02278$ $\omega_1 = 0.06398$ | Two-ratio vs. M0 | 10.94072 | 0.00094 |
|  | Free-ratio | -202667.8186 |  | Free-ratio vs. M0 | 2008.56299 | 0.00000 |
| ML tree | M0 | -203672.1001 | $\omega = 0.02272$ | | | |
| | Two-ratio | -203666.6298 | $\omega_0 = 0.02278$ $\omega_1 = 0.06398$ | Two-ratio vs. M0 | 10.94072 | 0.00094 |
|  | Free-ratio | -202747.0065 |  | Free-ratio vs. M0 | 1850.18729 | 0.00000 |
| Branch-site models |  |  |  |  |  |  |
| Bayesian tree | Null model | -202664.0952 | $P_0 = 0.77667$ $P_1 = 0.02562$ $P_{2a} = 0.19139$ $P_{2b} = 0.00631$<br>$\omega_0 = 0.02110$ $\omega_1 = 1.00000$ $\omega_{2a} = 1.00000$ $\omega_{2b} = 1.00000$ | | | |
| | Model A | -202663.7531 | $P_0 = 0.78464$ $P_1 = 0.02590$ $P_{2a} = 0.18340$ $P_{2b} = 0.00605$<br>$\omega_0 = 0.02115$ $\omega_1 = 1.00000$ $\omega_{2a} = 1.21998$ $\omega_{2b} = 1.21998$ | Model A vs. Null model | 0.68422 | 0.40814 |
| ML | Null model | -202664.0952 | $P_0 = 0.77667$ $P_1 = 0.02562$ $P_{2a} = 0.19140$ $P_{2b} = 0.00631$<br>$\omega_0 = 0.02110$ $\omega_1 = 1.00000$ $\omega_{2a} = 1.00000$ $\omega_{2b} = 1.00000$ | | | |
| | Model A | -202663.7531 | $P_0 = 0.77667$ $P_1 = 0.02562$ $P_{2a} = 0.19139$ $P_{2b} = 0.00631$<br>$\omega_0 = 0.02115$ $\omega_1 = 1.00000$ $\omega_{2a} = 1.22003$ $\omega_{2b} = 1.22003$ | Model A vs. Null model | 0.68422 | 0.40814 |

Moreover, many studies have shown that positive selection often occurs over a short period of evolutionary time and acts on only a few sites; thus, the signal for positive selection is usually swamped by those for continuous purifying selection that occur on most sites in a gene sequence [87,93]. In the present study, branch-site models were used to detect possible positively selected sites in the vesicomyid bivalves (Table 4). Ten residues, which were located in *cox1, cox3, cob, nad2, nad4* and *nad5*, were identified as positively selected sites with high BEB values (> 95%).

**Table 4.** Possible sites under positive selection in the Vesicomyidae lineage.

| Bayesian tree |  |  |  | ML tree |  |  |  |
| --- | --- | --- | --- | --- | --- | --- | --- |
| Gene | Codon | Amino acid | BEB values | Gene | Codon | Amino acid | BEB values |
| *cox1* | 529 | W | 0.967 | *cox1* | 529 | W | 0.967 |
| *cox3* | 998 | G | 0.956 | *cox3* | 998 | G | 0.956 |
|  | 1018 | W | 0.962 |  | 1018 | W | 0.962 |
|  | 1021 | T | 0.954 |  | 1021 | T | 0.954 |
| *cob* | 1131 | K | 0.982 | *cob* | 1131 | K | 0.982 |
|  | 1432 | A | 0.959 |  | 1432 | A | 0.959 |
| *nad2* | 2043 | K | 0.960 | *nad2* | 2043 | K | 0.960 |
| *nad4* | 2388 | E | 0.951 | *nad4* | 2388 | E | 0.951 |
| *nad5* | 2734 | P | 0.975 | *nad5* | 2734 | P | 0.975 |
|  | 2773 | S | 0.951 |  | 2773 | S | 0.951 |

It is well known that mitochondrial PCGs play a key role in the oxidative phosphorylation pathway; the above ten amino acid mutation sites are components of the respiratory chain and therefore may have important functions. As the first and the largest enzyme complex in the respiratory chain, the NADH dehydrogenase complex exercises the functions of proton pumps, and variation in loci may affect metabolic efficiency [90]. In this work, there were four positively selected sites located in the *nad2, nad4* and *nad5* genes. Similar results have been obtained in studies of the adaptive evolution of Tibetan horses, Chinese snub-nosed monkeys and Tibetan loaches, which live in high-altitude habitats [17,19,22]. Two residues in the *cob* gene were identified to be under positive selection. As a relatively conserved gene, *cob* plays a fundamental role in energy production in mitochondria. It catalyzes reversible electron transfer from ubiquinol to cytochrome *c* coupled to proton translocation [94]. Wide variation in the properties of amino acids was observed in functionally important regions of *cob* in species with more specialized metabolic requirements, such as adaptation to a low-energy diet or large body size and adaptation to unusual oxygen requirements or low-temperature environments [90,95]. Cytochrome c oxidase, which catalyzes the terminal reduction of oxygen and whose catalytic core is encoded by three mitochondrial protein-coding genes (*cox1, cox2* and *cox3*), has been proven to be a particularly important target of positive selection during hypoxia adaptation [96–97]. Four positively selected residues were detected in the *cox1* and *cox3* genes. For *C. marissinica*, functional modification mediated by positively selected mutations may increase the affinity between the enzyme and oxygen, thus allowing the efficient utilization of oxygen under hypoxia and maintaining essential metabolic levels.

The environment of deep-sea hydrothermal vents and cold seeps is characterized by darkness, a lack of photosynthesis-derived nutrients, high hydrostatic pressure, variable temperatures, low dissolved oxygen, and high concentrations of hydrogen sulfide (H_2_S), methane (CH_4_) and heavy metals, such as iron, copper and zinc. Previous studies have confirmed that all of the above environmental factors influence the process of mitochondrial aerobic respiration; for example, thirty potentially important adaptive residues were identified in the mitogenome of *S. leurokolos* and revealed the mitochondrial genetic basis of hydrothermal vent adaptation in alvinocaridid shrimp [65]. Similar findings have been reported in other deep-sea macrobenthos, such as the sea anemone *Bolocera* sp., starfish *Freyastera benthophila* and sea cucumber *Benthodytes marianensis* [64,98–99]. In the present study, ten potentially adaptive residues were identified in the *cox1, cox3, cob, nad2, nad4* and *nad5* genes, supporting the adaptive evolution of the mitogenome of *C. marissinica*. Our results at least partly explained how the deep-sea vesicomyid bivalves maintain aerobic respiration for sufficient energy in the extremely harsh deep-sea environment. The findings of this study could help deepen our understanding of the molecular mechanisms of adaptive evolution at the mitochondrial level in deep-sea organisms.

## Conclusion

This study characterized the complete mitogenome of the deep-sea vesicomyid bivalve *C. marissinica*, which is 17,374 bp in length and encodes 37 typical mitochondrial genes, including 13 PCGs, 2 rRNA genes, and 22 tRNA genes. All of these genes are encoded on the heavy strand. We analyzed the mitogenome organization, codon usage, control region features, gene arrangement, phylogenetic relationships and positive selection of *C. marissinica*. In the mitogenome of *C. marissinica*, tandem repeat sequences, “G(A)_n_T” motifs and AT-rich sequences were detected. In the family Vesicomyidae, we found that if the tRNA genes are not considered, the sequenced vesicomyid bivalves have a completely identical arrangement of PCGs. The phylogenetic analyses clustered *C. marissinica* with previously reported vesicomyid bivalves with high support values. Ten residues located in *cox1, cox3, cob, nad2, nad4* and *nad5* were inferred to be positively selected sites along the branches leading to vesicomyid bivalves, which may indicate that the genes were under positive selection pressure. This study probes the mitochondrial genetic basis of deep-sea adaptation in vesicomyids and provides valuable insight into the adaptation of organisms to the extreme deep-sea environment.

## Supporting information

**S1 Table. Species used for phylogenetic reconstructions.**

**S2 Table. Species used for CodeML analyses of selective pressure on mitochondrial genes.**

**S1 Fig. Putative secondary structures for the 22 transfer RNAs of the *C. marissinica* mitogenome.**

**S2 Fig. Phylogenetic trees derived from ML analyses based on nucleotide sequences of 9 mitochondrial protein-coding genes and 2 ribosomal RNA genes.**

**S3 Fig. Phylogenetic trees derived from Bayesian analyses based on amino acid sequences of 9 mitochondrial protein-coding genes and 2 ribosomal RNA genes.**

**S4 Fig. Phylogenetic trees derived from ML analyses based on amino acid sequences of 9 mitochondrial protein-coding genes and 2 ribosomal RNA genes.**

## Acknowledgments

The authors thank the captains and crews of the R/V Tan Suo Yi Hao and the pilots of the HOV “Shen Hai Yong Shi” for technical support. This research is funded by the National Key R&D Program of China (2018YFC0309804), the National Natural Science Foundation of China (NO. 41506173) and the Strategic Priority Research Program of the Chinese Academy of Sciences (No. XDB06010101).

